# Rap-protein paralogs of *B. thuringiensis*: a multifunctional and redundant regulatory repertoire for the control of collective functions

**DOI:** 10.1101/784611

**Authors:** Gabriela Gastélum, Mayra de la Torre, Jorge Rocha

**Author notes:** Address correspondence to Jorge Rocha. **Data Deposition Statement**: Analysis scripts and input files associated with reconstruction of the phylogenetic tree are available at https://github.com/gabyga16/rap_phylogenetics. A file with supplemental material is available.

## Abstract

Quorum Sensing (QS) are mechanisms of synthesis and detection of signaling molecules to regulate gene expression and coordinate behaviors in bacterial populations. In *Bacillus subtilis* (Bs), multiple paralog Rap-Phr QS systems (receptor-signaling peptide) are highly redundant and multifunctional, interconnecting the regulation of differentiation processes such as sporulation and competence. However, their functions in the *B. cereus* group are largely unknown. We evaluated the diversification of Rap-Phr systems in the *B. cereus* group as well as their functions, using *Bacillus thuringiensis* Bt8741 as model. Bt8741 codes for eight Rap-Phr systems; these were overexpressed to study their participation in sporulation, biofilm formation, extracellular proteolytic activity and spreading. Our results show that five Rap-Phr systems (RapC, K, F, I and RapLike) inhibit sporulation, two of which (RapK and RapF) probably dephosphorylate of Spo0F from the Spo0A phosphorelay; these two Rap proteins also inhibit biofilm formation. Five systems (RapC, F, F2, I1 and RapLike) decrease extracellular proteolytic activity; finally, four systems (RapC, F1, F2 and RapLike) participate in spreading inhibition. Our bioinformatic analyses showed that Rap proteins from the *B. cereus* group diversified into five pherogroups, and we foresee that functions performed by Rap proteins of Bt8741 could also be carried out by Rap homologs in other species within the group. These results indicate that Rap-Phr systems constitute a highly multifunctional and redundant regulatory repertoire that enables bacteria from the *B. cereus* group to efficiently regulate collective functions during the bacterial life cycle, in the face of changing environments.

**Importance:** The *Bacillus cereus* group of bacteria includes species of high economic, clinical, biological warfare and biotechnological interest, e.g. *B. anthracis* in bioterrorism, *B. cereus* in food intoxications and *B. thuringiensis* in biocontrol. Knowledge on the ecology of these bacteria is hindered due to our limited understanding about the regulatory circuits that control differentiation and specialization processes. Here, we uncover the participation of eight Rap quorum-sensing receptors in collective functions of *B. thuringiensis*. These proteins are highly multifunctional and redundant in their functions, linking ecologically relevant processes such as sporulation, biofilm formation, extracellular proteolytic activity, spreading, and probably other additional functions in species from the *B. cereus* group.

## Introduction

Bacteria perform many functions that depend on multicellular-like behaviors, such as cell differentiation and specialization. These behaviors, also known as collective functions, allow the emergence of complex ecological interactions, including cooperation and division of labor in biofilms (1, 2). Collective functions are only evident and effective when performed by large groups in bacterial populations or communities (3–6). Some of the most studied examples include bioluminescence by the squid symbiont *Vibrio fischeri* (7), or fruiting body formation during sporulation of *Myxococcus xanthus* (8).

In gram-positive bacteria, collective functions and the molecular mechanisms for their control have been widely studied in *Bacillus subtilis* (Bs). In Bs cultures, several mutually-exclusive cell-types have been identified (motile, competent, sporulating, cannibal, biofilm matrix producers, surfactant producers and miners (9, 10)), where emerging ecological interactions such as cooperation, cheating and cross-feeding, have been described (5, 6, 11). The presence of these cell differentiation phenomena and the resulting ecological interactions, ultimately affect the manifestation of collective traits such as sporulation efficiency, surface colonization, biofilm architecture complexity, etc. (2, 9, 12). These phenomena depend on global modifications of transcriptional regulation; they are triggered by environmental cues, stress conditions, cell-cell signaling, and are tightly modulated by complex, overlapping regulatory circuits (13–15).

Bacteria detect cell density through quorum sensing (QS), which depends on self-produced signaling molecules that accumulate in the extracellular space as the population grows. Specific receptors in the cell membrane or in the cytoplasm recognize these signaling molecules and regulate downstream cellular processes (16–18). Collective traits such as virulence, competence, sporulation and bioluminescence are regulated by QS. Gram-positive bacteria use small peptides as signaling molecules for QS (17).

The RRNPP family (Rgg, Rap, NprR, PlcR, PrgX) are intracellular QS receptors that regulate several functions across gram-positive bacteria (19–21). Genes coding for receptor proteins and their associated signaling peptides are encoded in transcriptional cassettes (22). Rgg, NprR, PlcR and PrgX proteins are transcriptional activators that bind directly to DNA in quorum state. Rap proteins, however, lack a DNA binding domain and they function by binding and inhibiting proteins, specifically response regulators and transcriptional activators (21, 23, 24). Twelve Rap paralogs (RapA, B, C, D, E, F, G, H, I, J, K, 60) control diverse functions in *B. subtilis* 168 (Bs168). The RapG-PhrG pair regulates the activation of DegU, a transcriptional regulator that controls *aprE* and *comK* genes encoding for extracellular proteases and a transcription factor for competence in Bs, respectively (15, 25); ComA – the master regulator of competence genes – is repressed by RapC, D, F, G, H, K and Rap60 (14, 26–31); Spo0A – the transcriptional activator of many differentiation genes – is indirectly regulated by RapA, B, E, H, J, and Rap60 (24, 31–35). Hence, Rap protein paralogs from Bs are highly multifunctional and redundant and they connect several differentiation processes and coordinate collective traits.

Spo0A is activated by phosphorylation through a multicomponent phosphorelay system. Up to five kinases auto-phosphorylate in response to intracellular and environmental stress signals and transfer the phosphate group to Spo0F, which is then transferred to Spo0B and finally to Spo0A (36). Spo0A-P activates the transcription of multiple genes, including biofilm formation (at low concentrations) and early sporulation genes (at high concentrations (13)). Rap QS proteins prevent the phosphate transfer in the phosphorelay by binding to Spo0F (32, 37).

While the regulation of collective traits in Bs is well known, these phenomena remain largely understudied in the *B. cereus* group, which includes bacteria with clinical and biotechnological relevance (38). Although Bs and *B. cereus* group species share similar characteristics such as the sporulation process, the Spo0A phosphorelay components, and have many protein families in common, they also present notorious genetic differences (39). In *B. thuringiensis* (Bt, the most widely used biopesticide), the Spo0A phosphorelay is modulated by the bifunctional QS receptor NprR, which is not present in Bs (40–42). On the other hand, ComA and DegU response regulators are not encoded in Bt. Additionally, Rap-Phr QS systems also differ in both groups. These QS systems have evolved by duplication and divergence mechanisms; even though multiple Rap proteins paralogs are also found *B. cereus* group species, they have evolved independently and no Rap homologs are shared between the two groups (43, 44). Therefore, it is not possible to predict the functions of Rap proteins in the *B. cereus* group based on what is known of Rap proteins from Bs.

Some Rap-Phr systems from species of the *B. cereus* group have been studied. First, Rap BXA0205 and BA3790 from *B. anthracis* str. A2012, were demonstrated to regulate sporulation initiation and to dephosphorylate Spo0F (45). Later, it was shown that Rap8 from Bt-HD73, regulates the sporulation and biofilm formation processes *in vitro* (46). A more recent study showed the participation of Rap6, 7 and 8 – also known as RapC, K and RapF, respectively (47) – in the modulation of the sporulation process in Bt407 (48). However, other Rap paralogs with unknown functions have been identified in the genomes of *B. cereus* group bacteria (44, 47) that may be relevant to their ecology.

In this study we aimed at evaluating the diversification of the Rap-Phr systems in the *B. cereus* group as well as their functions, using *Bacillus thuringiensis* Bt8741 as model. We generated eight Rap-overexpression strains of Bt8741 to evaluate the role of each Rap paralog in sporulation efficiency, biofilm formation, extracellular proteolytic activity and spreading. We also studied the evolution of Rap-Phr paralogs in the *B. cereus* group, by identifying Rap homologs from other species and analyzing its phylogeny. This allowed the prediction of their functions, based on those of Rap proteins from Bt8741.

## Results

### Spo0F-binding residues from Bs-RapH are conserved in Rap proteins from Bt8741

In order to predict the capacity of Rap proteins from Bt407 (a strain closely related to Bt8741) to bind to Spo0F, we analyzed the conservation of the amino acids previously reported to be involved in Spo0F-binding by RapH from *B. subtilis*. In this analysis, we included reported sequences of Bs-Rap proteins that bind to Spo0F (RapA, B, E, H, J) as well as the sequence of RapD from Bs, which does not bind to Spo0F (34) (Fig. 1). We found more conservation of the functional amino acids of RapH, in the sequences of both Bs168 and Bt407, compared to the corresponding full sequences (Fig. 1B). In Bs168, the full sequence conservation of the Rap proteins known to bind to Spo0F (RapA, B, E, J) compared to RapH, ranged from 59% to 66%, and the functional amino acids conservation percentage, from 82.3% to 100%. In RapD, which does not bind to Spo0F, the full-length sequence is conserved at 50% and the functional residues are only 64.7% conserved (Fig. 1B). In the case of Rap proteins from Bt407, the full sequence conservation in comparison to RapH of Bs168 ranged from 45% to 48%. On the other hand, conservation of the functional residues ranged from 64.7% to 88.2% (Fig 1B). Since more conservation occurs in the Spo0F-binding functional residues, these residues could be important for the function of Bt Rap proteins.

**Figure 1.**
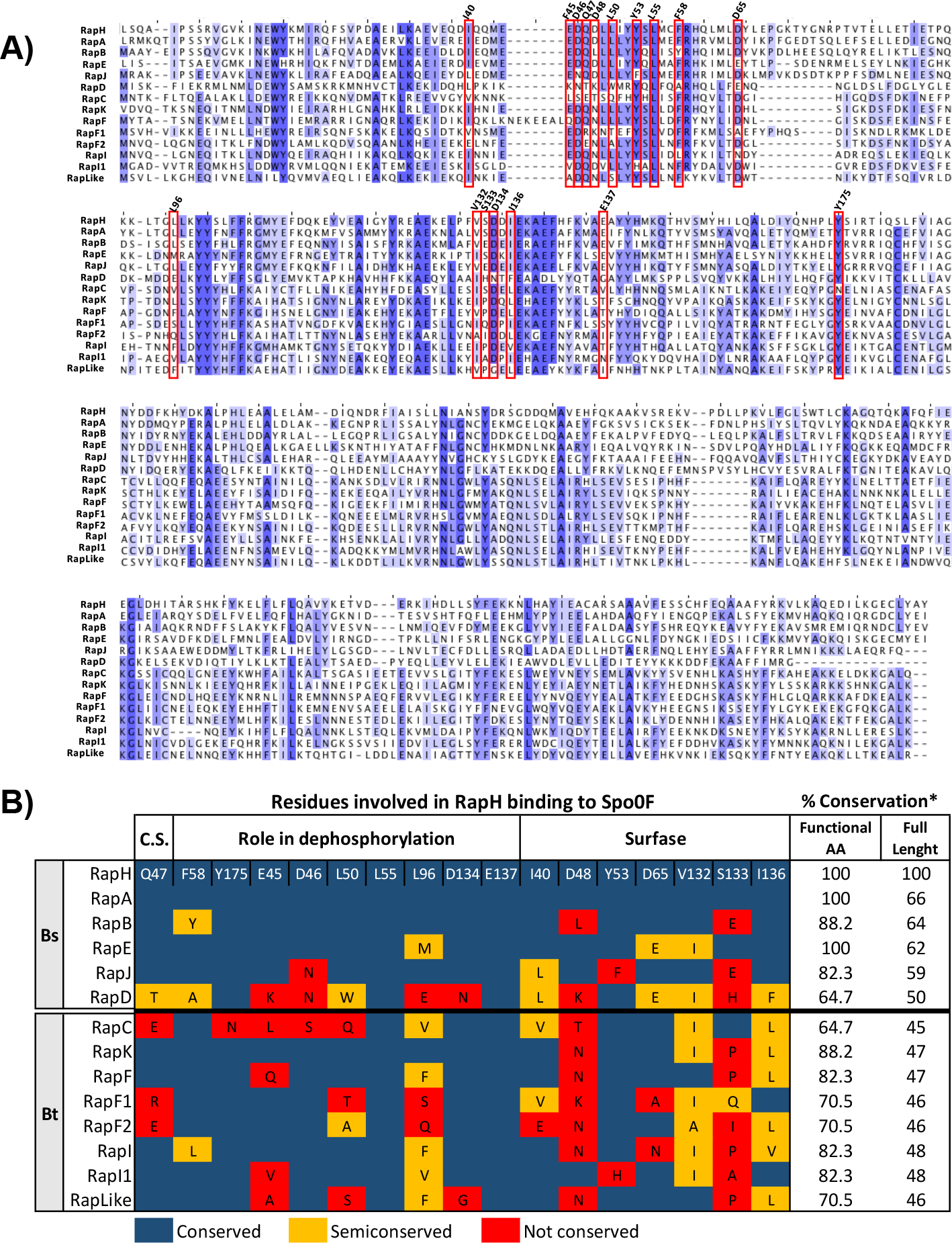
Prediction of the capacity of Rap proteins from Bt8741 to bind and dephosphorylate Spo0F. A) Multiple sequence alignment of the complete amino acid sequences of RapH, A, B, E, J and RapD from *B. subtilis* 168, and eight Rap proteins from Bt8741. Blue highlights indicate highly conserved amino acids. Residues involved in the RapH-Spo0F binding are indicated in red rectangles and its position in RapH is shown on top of the alignment. B) Conservation of residues involved in RapH binding to Spo0F. Residues were considered as semiconserved when a functional amino acid of RapH was substituted with another amino acid with similar characteristics. Bs, *Bacillus subtilis* 168; Bt, *Bacillus thuringiensis* 407; C.S., Catalytic Site; *Percentage of conserved and semiconserved amino acids in pairwise alignment to RapH.

RapK exhibited the highest conservation percentage of Spo0F-binding residues (88.2%), followed by RapF, I and RapI1 (82.3%), RapF1, F2 and RapLike (70.5%) and finally RapC, with 64.6%. Although RapF1 and RapF2 had a high conservation of functional residues, neither these Rap paralogs, nor RapC, conserve the residue Q47 found in the catalytic site and previously shown to be essential for the phosphatase activity of RapH (34). This analysis enables the prediction that some Rap protein paralogs from Bt8741, with a high conservation percentage of putative Spo0F-binding amino acids, could dephosphorylate Spo0F, while other paralogs could have evolved to participate in other regulatory processes. Indeed, RapK, RapF and unexpectedly RapC from Bt407, Rap8 from Bt-HD73 (ortholog to RapI from Bt407) and Rap BXA0205 and BA3790 from *B. anthracis*, (homologs of RapK and RapF2, respectively) have been shown to participate in the modulation of sporulation (45, 46, 48). Previous to this work, RapF1, I1, and RapLike from Bt407 (or its homologs in other species), had not been tested for their role in sporulation.

### RapC, K, F and RapLike control sporulation in Bt8741

We constructed nine Rap-overexpression strains in the Bt8741 background (Table S1), one for each endogenous Rap protein identified in Bt407 (RapC, K, F, F1, F2, I, I1, Like) and one more for RapA from Bs168 (RapA_Bs_). We also generated a control strain of Bt8741 carrying the empty plasmid pHT315-P_*xylA*_ (Table S1). DNA sequencing showed correct, in-frame insertion of P_*xylA*_ and *rap* genes in the pHT315 plasmid (not shown). We followed a growth time-course experiment of all strains in shaking flasks for 24 hours and confirmed that neither xylose addition, nor Rap overexpression, affected bacterial growth (Fig. S1).

In order to identify the Rap proteins involved in the regulation of sporulation initiation, we studied the effect of Rap overexpression in the sporulation efficiency of each strain. In this experiment, we observed that both addition of xylose to the culture medium and the presence of *rap* genes in the plasmid had minor effects on total and thermoresistant CFU counts of Bt8741 at 72 h. In the control strain, addition of xylose caused a decrease of ≈1 log10 in total and thermoresistant CFU (Fig. S2A and S2B). Similarly, when *rapF1* and *rapF2* genes were carried in the plasmid – but not overexpressed – total CFU decreased by up to one logarithm of CFU in comparison to the control strain (Fig. S2A). Additionally, sporulation decreased one logarithm in strains carrying *rapF1*, *rapF2* and *rapI1* in comparison to the control strain when overexpression was not induced (Fig. S2C). These unspecific effects were probably related to basal expression from the P_*xylA*_ promoter, even when xylose is not added, since pHT315 replicates at 15 copies per cell (49). However, the most dramatic effect was found in thermoresistant CFU of strains overexpressing Rap proteins (Fig. S2D).

In spite of the unspecific effect of xylose addition on growth and sporulation, sporulation efficiency of the control strain remained unchanged by the addition of inducer (Fig. 2). In contrast, overexpression of RapA_Bs_ caused a decrease in sporulation efficiency from 7.9% to 0.0005%. In fact, thermoresistant CFU were undetectable when RapA_Bs_ was overexpressed (Fig. S2D). We also found undetectable levels of spores in strains overexpressing RapK and RapF (Fig. S2D). Sporulation efficiency decreased from 32.93% to 0.0002% in the strain overexpressing RapK and from 9.24% to 0.0026% in the strain overexpressing RapF (Fig. 2). In Bs, RapA dephosphorylates Spo0F in the Spo0A phosphorelay (32) and this result indicates that it performs the same function in B8741; furthermore, it suggests that RapK and RapF carry out the same mechanism for regulation of sporulation initiation.

**Figure 2.**
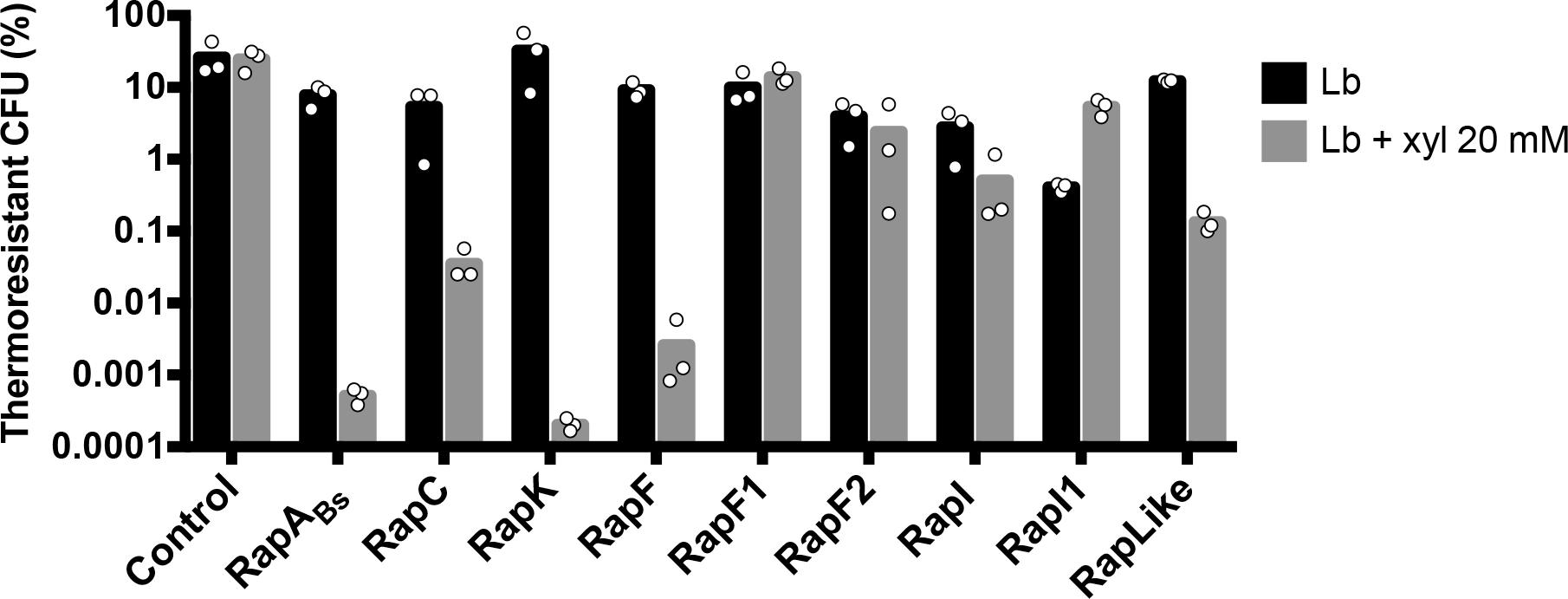
Sporulation efficiency of Bt8741 carrying overexpression plasmids for Rap proteins, with and without addition of inducer. In the cases where thermoresistant CFU were undetectable, we considered a value of 166 spores/ml, which is the detection limit for this assay. Columns represent average of three individual measurements, shown as dots.

Strains carrying P_*xylA*_*’rapC* and P_*xylA*_*’rapLike* also exhibited reduced sporulation efficiency when xylose was added to the medium. Sporulation efficiency decreased from 5.43% to 0.0357% and from 12.34% to 0.1352% when RapC and RapLike were overexpressed, respectively (Fig 2). Additionally, RapI overexpression slightly decreased sporulation efficiency, from 2.82% to 0.51%. Sporulation efficiency was not decreased by the overexpression of either RapF1, F2, I or RapI1.

Samples of the Rap-overexpressing strain cultures at 72 h were observed in a microscope. We detected free spores and bacterial debris in all cultures, when Rap proteins were not overexpressed (Fig. S3). Figure 3 shows representative fields of view with cells from each induced culture. Strains overexpressing Rap proteins that did not affect sporulation efficiency (RapF1, F2 I1) showed cell morphology similar to that of the control strain, i.e. a sporulated bacilli with defined endospores. In samples from strains overexpressing RapA_Bs_, K and RapF, that had acutely decreased sporulation efficiency, we observed chained, wrinkled cells with no spores (Fig. 3). On the other hand, cells from strains overexpressing RapC and RapLike, were observed as rod-shaped and no spores were evident (Fig. 3). Finally, in cells from the strain overexpressing RapI, which had a slight effect on sporulation efficiency, cell morphology was similar to strains overexpressing RapF1, F2, I1 and the control strain (Fig. 3), showing a defined endospore.

**Figure 3.**
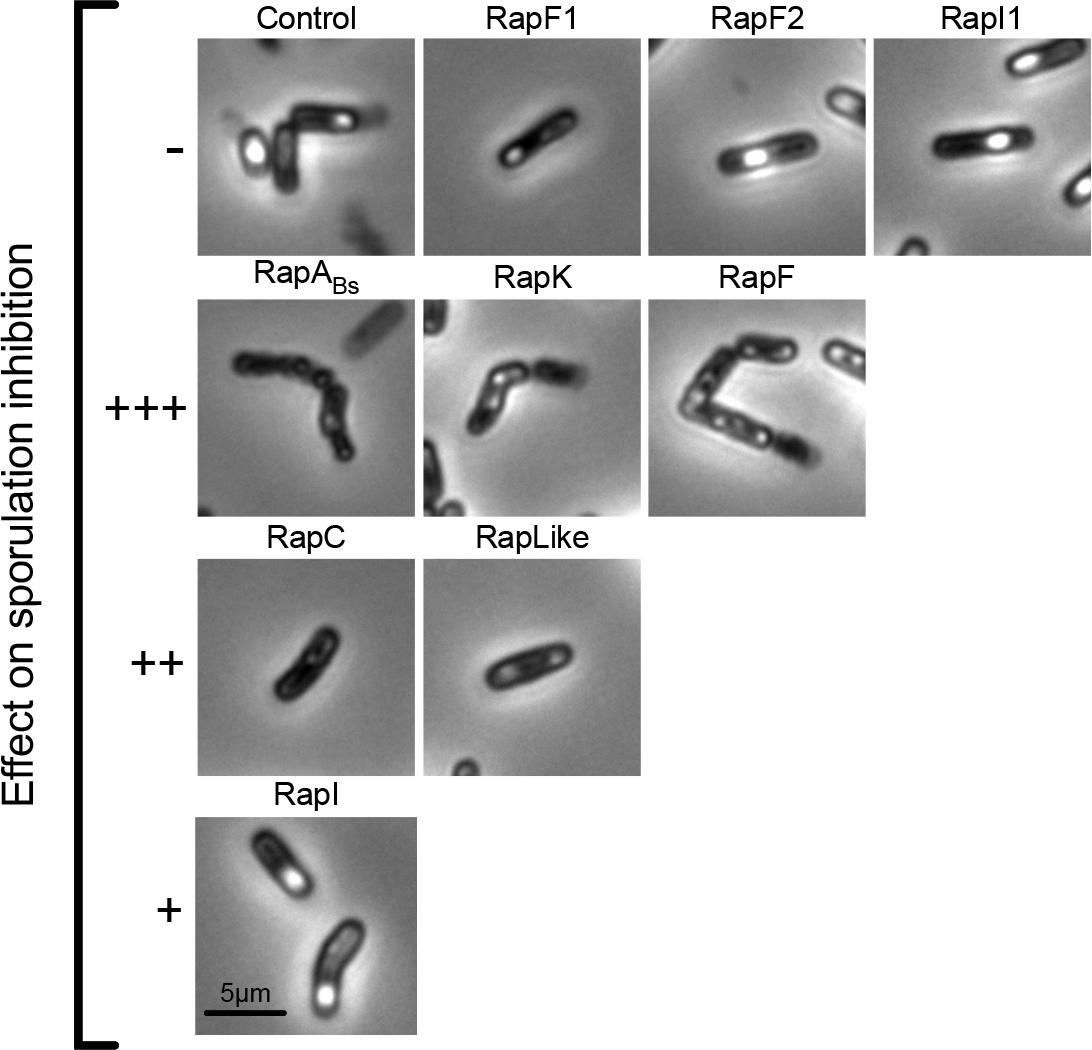
Cell morphology of strains with induced Rap protein overexpression at 72 h. Phase contrast microscopy 63× and 1.8× magnification. −, no effect; +, decrease under 10-fold; decrease between 90 and 160-fold; +++, decrease >1,000-fold.

### Overexpression of RapF and RapK prevents biofilm formation of Bt8741

In nature, over 80% of bacteria live in biofilms (49), therefore, biofilm formation is likely a relevant trait – albeit an understudied one – during the life cycle of Bt and other bacteria belonging to the *B. cereus* group. To determine which Rap proteins were involved in the regulation of biofilm development in Bt8741, we quantified biofilm formation of the Rap-overexpression strains in the air-liquid interphase at 48 h. For this, we suspended the cells form the biofilm and measured optical density (OD_600_). Since 20 mM of xylose in the media caused a complete inhibition of biofilm formation in the Bt8741 control strain (not shown), we first tested the effect of xylose concentration on this phenotype. We found that biofilm formation was not affected at 2 mM, but was decreased at higher concentrations of 5, 10 and 15 mM (Fig. S4); therefore, overexpression of Rap proteins was performed with 2 mM of xylose (50).

Overexpression of RapK and RapF caused an inhibition of biofilm formation of Bt8741 (Fig. 4A), evident by the significant decrease (p<0.0001) in the OD_600_ measured from a sample obtained from the surface of the culture (Fig. 4B). The OD_600_ of the biofilms decreased from 0.7115 to 0.0977 and from 0.6577 to 0.0961 in strains overexpressing RapK and RapF, respectively (Fig. 4B). On the other hand, biofilms were normally formed by strains overexpressing RapA_Bs_, C, F2, I, I1 and RapLike (Fig. 4). Interestingly, the strain overexpressing RapF1 was unable to form biofilms even when P_*xylA*_*’rapF1* was not induced (Fig. 4A and B).

**Figure 4.**
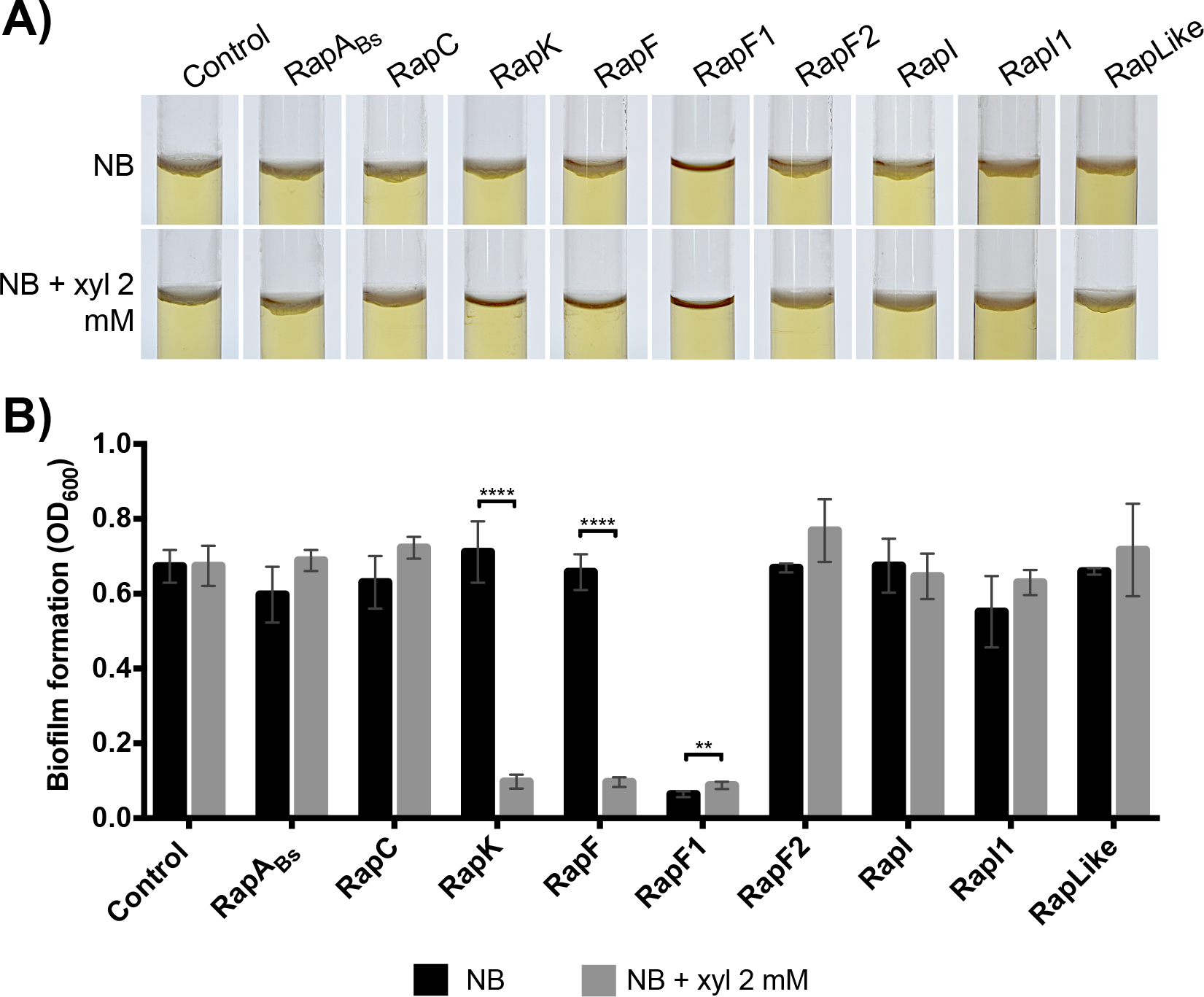
Biofilm formation of Rap-overexpression strains at 48 h. A) Biofilms formed in the liquid-air interphase in 13 × 100 mm glass tubes at 48 h. Biofilms are identified as a white layer on the surface. B) Biofilm formation quantification of Rap-overexpression strains in induced and not induced media after 48 h. Columns represent average of 5 replicates ± SD. NB, Nutrient Broth; **, p<0.005; ****, p<0.0001.

In order to discard possible global growth defects in this assay when RapK and RapF were overexpressed, we measured planktonic growth through OD_600_ of the liquid culture media from the same experiments where biofilm formation was assessed. We found that planktonic growth was higher in conditions where a biofilm was not formed (Fig. S5). This suggests that RapK and RapF specifically inhibit biofilm formation (e.g. secretion of extracellular matrix components).

### Extracellular proteolytic activity is downregulated by RapC, F, F2, I1 and RapLike in Bt8741

In Bt, the production of extracellular proteases is crucial during its necrotrophic phase, i.e. development in insect cadavers. We tested the role of Rap proteins in extracellular proteolytic activity by measuring the effect of Rap overexpression on hydrolysis halos of colonies on milk agar (MA) plates. Addition of xylose in the media had no effect (p>0.05) on the hydrolysis halo of the control strain (Fig. 5). In contrast, overexpression of RapC, F, F2, I1 and RapLike decreased the halo area (p<0.05; Fig. 5B). In these strains, the halo area decreased to 41.98%, 37.81%, 46.65%, 47.51% and 34.93%, respectively (Fig. 5B, Fig. S6) compared to the halos in plates where overexpression was not induced (100%). Proteolytic activity of strains overexpressing RapA_Bs_, K, I and RapF1 was not affected by the induction (p>0.05; Fig. 5A and B).

**Figure 5.**
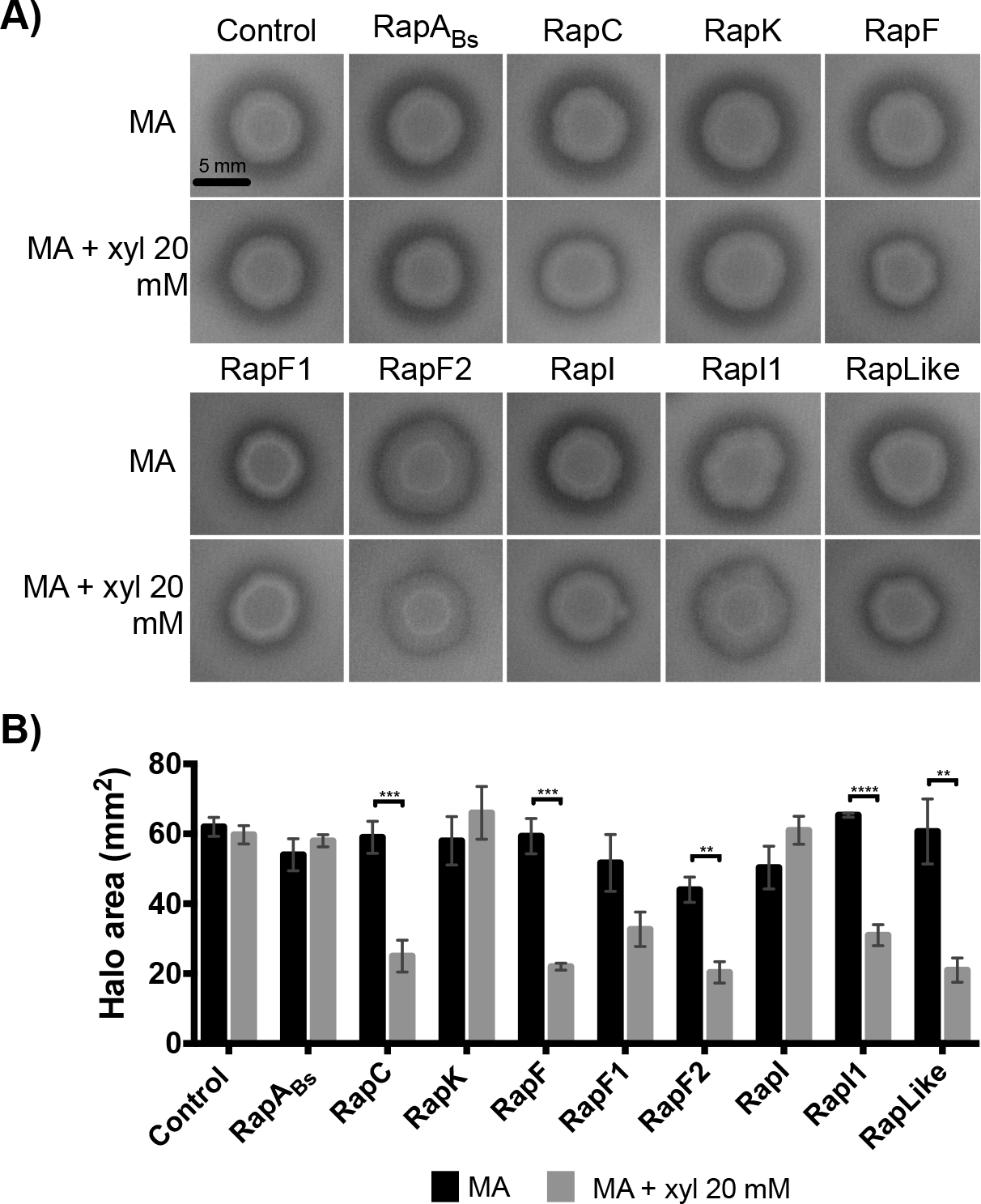
Extracellular proteolytic activity of Rap-overexpression strains. A) Effect of Rap protein overexpression in the hydrolysis halo of Rap-overexpression strains colonies. B) Hydrolysis halo area with and without Rap-overexpression induction. Columns represent average of 3 replicates ± SD. MA, milk agar; **, p<0.005; ***, p<0.0005; ****, p<0.0001.

### RapC, F1, F2 and RapLike regulate spreading of Bt8741 colonies

Colonies of Bt8741 present a spreading phenotype that could be associated to its capacity to colonize hosts and habitats. Similar passive motility phenotypes have been described in other species of *Bacillus*, associated to the production of extracellular surfactant molecules (51–53). To gain insights on this understudied collective trait, we determined the effect of Rap protein overexpression on radial spreading of colonies of Bt8741 growing on agar media.

We observed that addition of xylose in the media did not affect the spreading of the control strain (Fig. 6). In contrast, the overexpression of RapC, F1, F2 and RapLike caused a decrease in spreading (p<0.05) of Bt8741 colonies at day 7 (Fig. 6A and B). The overexpression of RapC reduced the colony dispersion from 5.15 mm to 0.49 mm (reduction of 90.4%); RapF1, from 3.73 mm to 1.83 mm (decrease of 50.9%); RapF2 from 5.05 mm to 0.78 mm (decrease of 84.5%); and RapLike from 3.64 mm to 0.65 mm (decrease of 82.1%) (Fig. 6B). Spreading inhibition is evident in the colony morphology of these strains (Fig. 6C). We observed that the overexpression of RapC, F2 and RapLike, completely eliminated this phenotype, while overexpression of RapF1 only decreased spreading (p<0.05) (Fig. 6B and C).

**Figure 6.**
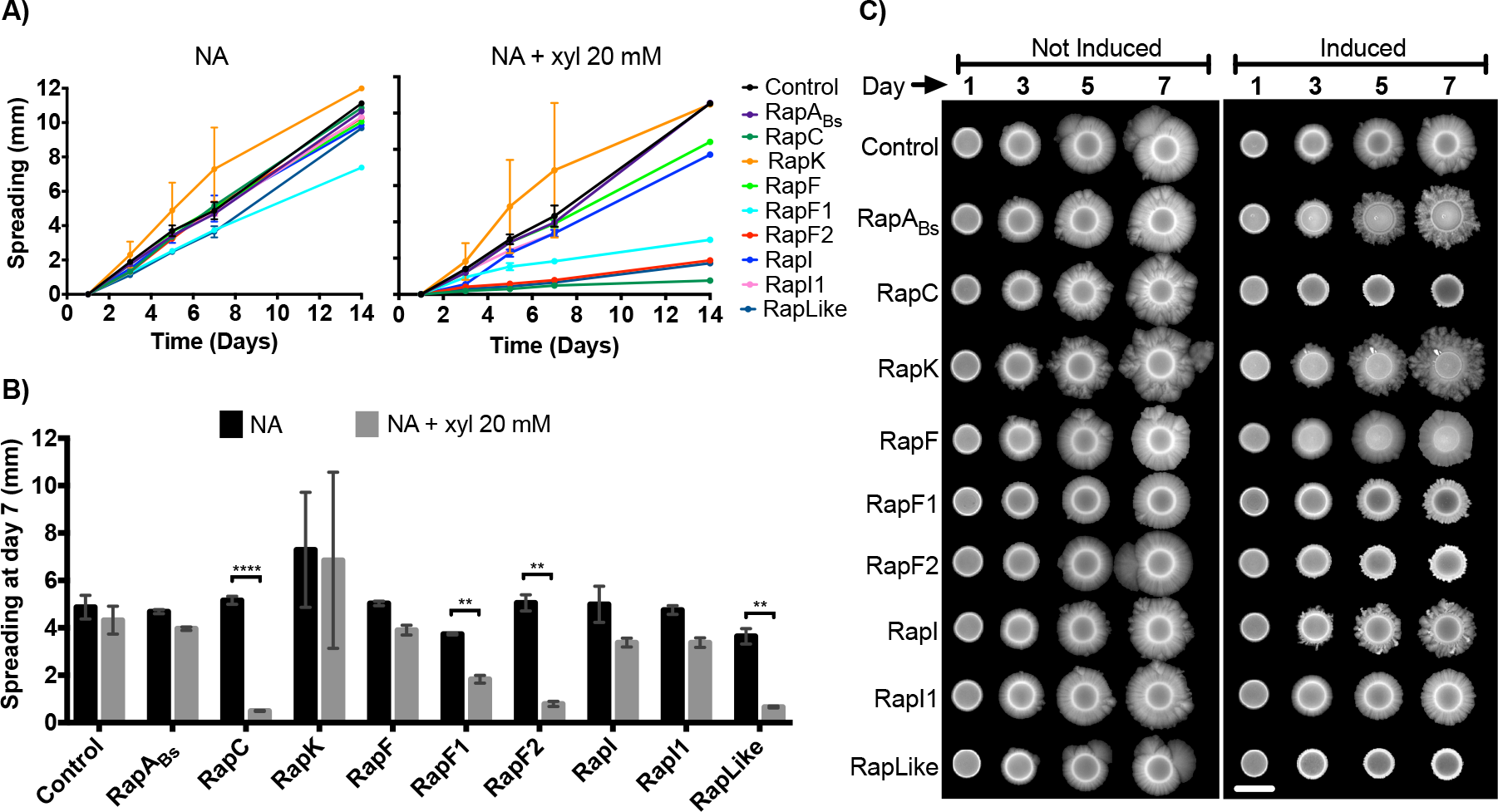
Spreading phenotype of Rap-overexpression strains. A) Spreading kinetics of Rap-overexpression colonies on agar. Each point represents the media of triplicates ± SD; only one data point is shown at day 14. B) Spreading quantification of Rap-overexpression colonies at day 7. Columns represent average of triplicates ± SD. C) Pictures of representative Rap-overexpression strains spreading during 7 days. Scale bar indicates 5 mm. NA, Nutrient Agar; **, p<0.005; ****, p<0.0001.

The overexpression of RapA_Bs_, K, F, I and RapI1 did not affect the spreading of Bt8741 (p>0.05) (Fig. 6B). Spreading of the strains carrying overexpression plasmids for these Rap proteins ranged from 4.68 mm to 7.29 mm without induction and from 3.37 mm to 6.84 mm when induced (Fig. 6B). In some cases, Rap overexpression affected colony morphology, i.e. colonies of strains overexpressing RapABs, RapK and RapI showed an increased dendritic phenotype; however, the spreading phenotype measured as colony radius, was still present (Fig. 6C).

### Rap-Phr systems diversified into five pherogroups in the *B. cereus* group

In order to predict the functions of Rap paralogs in *B. cereus* (Bc), *B. anthracis* (Ba), *B. mycoides* (Bm), *B. pseudomycoides* (Bps) and *B. cytotoxicus* (Bcyt), we analyzed their sequences to deduce the evolution of Rap proteins in these species and identify their putative signaling peptide sequences (mature Phr). Additional to the 8 *rap* genes in Bt407 (of which 4 are located in the chromosome, and 4 in plasmids) we found 32 *rap-phr* systems in the *B. cereus* group (Table S2), 30 of which are located in the chromosome and 2 in plasmids (Table 1).

**Table 1.** Rap-Phr systems encoded in *B. subtilis* 168 and in species from the *B. cereus* group.

| Specie | Number of Rap-Phr systems | Location |  |
| --- | --- | --- | --- |
|  |  | Chromosome | Plasmid |
| *Bacillus subtilis* 168 | 12 | 11 | 1 |
| *Bacillus cereus* ATCC 14579 | 5 | 5 | 0 |
| *Bacillus anthracis* str. A0248 | 6 | 5 | 1 |
| *Bacillus thuringiensis* 407 | 8 | 4 | 4 |
| *Bacillus mycoides* ATCC 6442 | 5 | 4 | 1 |
| *Bacillus pseudomycoides* DMS 12442 | 8 | 8 | 0 |
| *Bacillus cytotoxicus* NVH 391-98 | 8 | 8 | 0 |

The phylogeny of Rap proteins from the *B. cereus* group shows that clades are composed of Rap proteins from different species, i.e., phylogenetically close Rap homologs can be found in different species. This suggests that Rap-Phr divergence occurred before speciation in this group (Fig. 7). Hence, it is possible that Rap functions uncovered in this work could be extrapolated to the rest of the *B. cereus* group, e.g. Rap proteins found in the same clade as BtRapK and BtRapF (Bps28285, Bps05775, Bps24285, Bcyt05320, Bc1026, Ba05875, Ba29315, Bcyt11595, Bcyt05405 and Bcyt02700) may modulate sporulation initiation and biofilm formation. Since we found that several Rap paralogs are coded in every species of the *B. cereus* group, we suggest that they could regulate a variety of collective functions in all these species, as we describe here for Bt8741.

**Figure 7.**
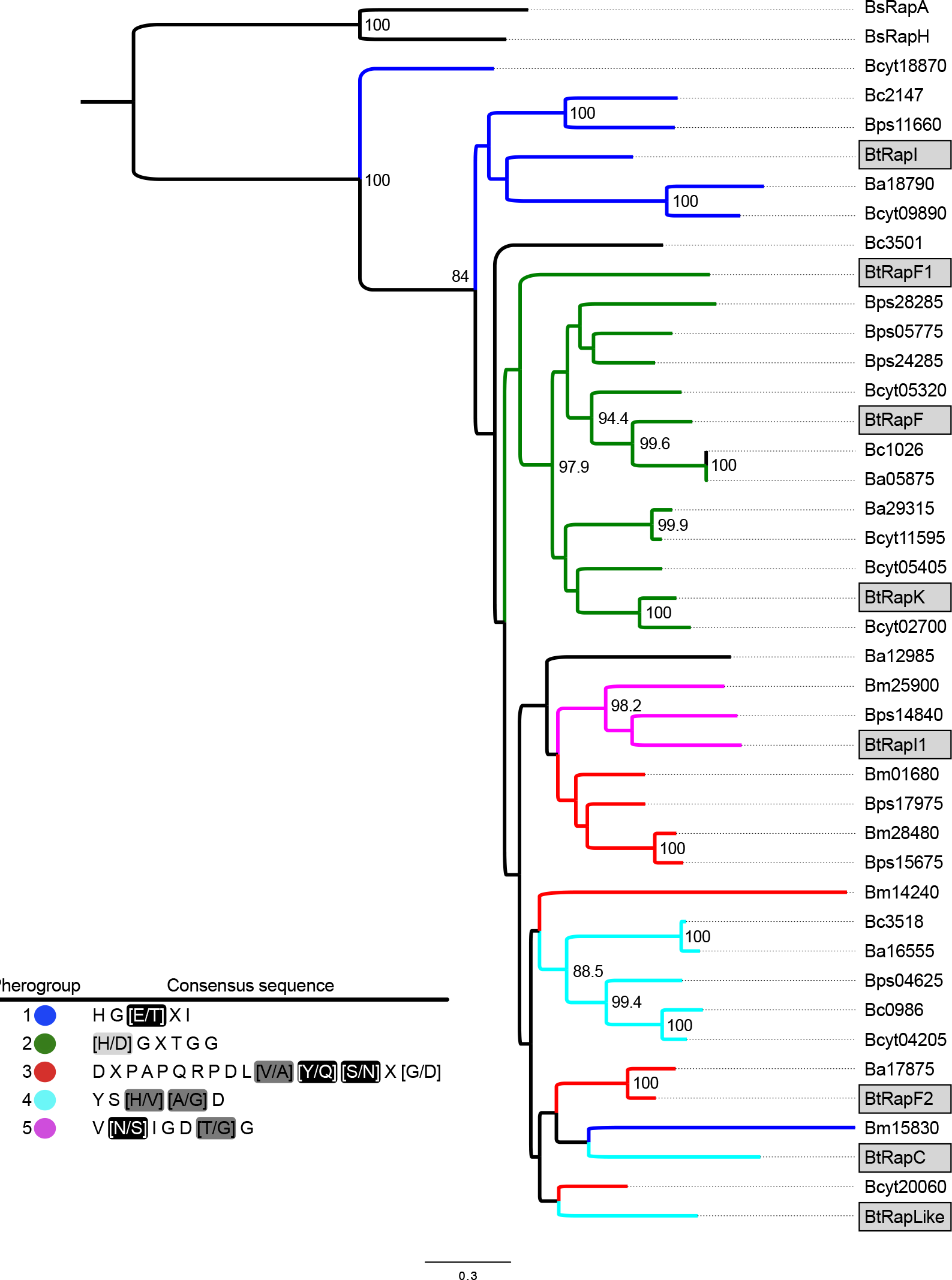
Maximum likelihood phylogeny of Rap proteins from the *B. cereus* group. Rap proteins from Bt8741 are highlighted in gray boxes. Branches from each pherogroup are identified in colors. Bootstraps higher than 80% are shown in each node. Insert table: pherogroups and consensus sequences of mature Phr. Semiconservations in the consensus sequences are highlighted: black, polar residues; gray, hydrophobic residues; silver, polar and charged residues.

We identified 5 pherogroups, each with a putative mature Phr peptide consensus sequence. All five pherogroups include Rap proteins from different species (Fig. S7). These five pherogroups are identified with colors in the branches of the phylogeny in figure 7. We found that the mature Phr corresponding to pherogroups 1 and 2 are located at the C-terminal domain of the pro-peptides (exported Phr sequence). RapI, F, F1 and RapK from Bt are found at these pherogroups. On the other hand, for pherogroup 3 – where RapF2 from Bt is found – consensus sequences are located at the N-terminal domain of the pro-peptide. Finally, putative mature Phr peptides from pherogroups 4 and 5 – which include RapLike, C and RapI1 from Bt – are located in the middle of the exported sequence (Table S3). We observed that only Bt and Bps encode Rap proteins from all five pherogroups; Rap proteins from Ba and Bcyt are found in pherogroups 1, 2, 3 and 4; Rap proteins in Bm correspond to pherogroups 1, 3 and 5; Rap proteins from Bc are found only in pherogroups 1 and 4 (Fig. 7). Pherogroup 1 is the only one present in all evaluated species.

## Discussion

Few studies have addressed multicellular behaviors such as differentiation, cell-specialization, collective functions, and the resulting ecological interactions in species from the *B.* cereus group (11, 54). Similarly, molecular mechanisms for the control of differentiation processes in the *B. cereus* group bacteria remain understudied (45, 46, 48, 54, 55). Here we show that Rap-Phr systems in Bt8741 regulate collective functions such as sporulation, biofilm formation, production of extracellular proteases and spreading motility. In fact, Rap-Phr systems in this strain are highly multifunctional and redundant, since five out of eight Rap paralogs modulate more than one collective trait, and all four collective traits studied were inhibited by more than one Rap protein. Hence, Rap paralogs appear to constitute a regulatory repertoire that allows Bt populations to respond efficiently to environmental changes, which contributes to fitness of the population.

Although it is well known how the Rap-Phr systems participate in differentiation processes of the gram-positive model bacteria Bs, speciation resulted in divergent Rap proteins in the *B. cereus* and *B. subtilis* groups (43, 44). Therefore, no homologs are shared between the groups; however, in both cases, speciation resulted in the presence of multiple Rap paralogs per genome. We propose that Rap proteins that are phylogenetically close to Rap proteins from Bt8741, could have the same functions in other bacteria of the *B. cereus* group.

It is not yet clear how bacteria benefit from keeping multiple receptor-signaling peptide gene pairs comprising this complex signaling network of Rap-Phr systems; however, it has been shown that redundancy in Rap-Phr systems in Bs has been selected for because it provides social advantages (56). Because Rap proteins have a repressive function upon its target, the gain of a novel Rap-Phr system for the regulation of extracellular public good production enables a facultative cheating mechanism in which variants with an extra system exploit their ancestral strain. Here we showed that extracellular public goods such as biofilm matrix components, extracellular proteases or surfactants, are likely controlled by Rap proteins in Bt8741; therefore, the same facultative cheating mechanism could be expected during duplication of *rap-phr* genes in the *B. cereus* group. This represents a selective advantage by a fitness increase of the novel population. Multifunctionality seems to have evolutionary advantages as well. Perhaps, because Rap-Phr systems are known to be parallel signaling pathways (44) they are not all activated simultaneously; instead, some of them may be active only under specific conditions, achieving the regulation of various differentiation processes and collective functions while optimizing energetic costs. Overall, keeping multiple redundant and multifunctional Rap paralogs that control important collective functions results in a better adaptation and population survival in nature.

Sporulation in the *Bacillus* genus is essential for bacterial survival and dissemination in their habitats; it is also important for the biotechnological uses of *Bacillus* species. Six Rap-Phr systems from Bs, including RapA, negatively regulate Spo0A phosphorelay by dephosphorylating Spo0F, and therefore prevent the activation of Spo0A (32). We found that RapA_Bs_, retained this function when it was overexpressed in Bt8741. Furthermore, five Rap-Phr systems from Bt8741 (RapK, F, C, Like and RapI) also regulate sporulation in this species. We propose that RapK and RapF may function by dephosphorylating Spo0F, similar to the mechanism carried out by RapA in Bs. This suggestion is supported by three findings: 1) both RapK and RapF retain the highest conservation of Spo0F binding residues from RapH, including the catalytic residue Q47; 2) their overexpression resulted in undetectable number of spores, similar to RapA_Bs_ overexpression; 3) the overexpression of RapA_Bs_, K and RapF caused an identical cell morphology in the three overexpressing strains. Additionally, RapK and RapF are closely related and both belong to pherogroup 2, which may indicate that they resulted from a gene duplication event of a Rap ancestor that dephosphorylated Spo0F. Other Rap proteins that decreased sporulation efficiency are RapC, Like and RapI; of these, RapC does not contain the catalytic site residue Q47. Further studies are needed in order to elucidate the mechanisms by which all these receptors regulate sporulation in Bt and other species from the *B. cereus* group.

RapK and RapF are the only Rap proteins from Bt8741 that prevented biofilm formation. Because Spo0A-P levels regulate both sporulation and biofilm formation in Bs, we speculate that bifunctionality of RapK and RapF in Bt8741 results from their activity on Spo0F. We noted, however, that the overexpression of RapA_Bs_, which completely prevented sporulation, did not affected biofilm formation in Bt. Overexpression of RapA_Bs_ may allow low levels of Spo0A-P in Bt8741, which in Bs are sufficient for the activation of genes related to production of extracellular matrix components, but not for the activation of early sporulation genes (13). This picture is probably more complex, as different feedback loops modulate sporulation and biofilm formation (57). It is noteworthy that overexpression of RapA_Bs_, in Bt8741 did not inhibit any phenotype, other than sporulation; this reflects the fact that Rap proteins co-evolve with specific protein targets in each bacterial species; it also indicates that Rap target regulators involved in the control of extracellular proteases and spreading may not be conserved betweem Bs and the *B. cereus* group.

We suggest that Rap proteins have diversified according to the ecological needs of each species. For example, Bs is a soil dwelling bacteria found associated to rhizosphere forming biofilms (58). In Bs, six Rap proteins modulate Spo0A-P levels (21, 59), affecting sporulation and biofilm formation. Here we demonstrate that five Rap proteins modulated sporulation (RapC, K, F, I and RapLike) while only two of these (RapK and RapF) affected biofilm formation, perhaps through the Spo0A phosphorelay. This highlights the importance of sporulation regulation in both species and that probably, biofilm formation is not as essencial in the lifecylce of Bt, as it is in Bs. In contrast, Bt is a soil inhabitant, insect patogenic and necrotrophic bacteria (60). In this species, extracellular protease production is essential for nutrient scavenging, which is normally associated to the necrotrophic stage of bacterial development in the insect cadaver (40). Additionally, it could be relevant during the transition from exponential growth to stationary phase in controlled fermentations or for adaptation against fluctuations in nutrient availability in the environment. While only one of the twelve Rap proteins from Bs modulates its extracellular proteolytic activity (RapG) (25), Bt has extended the modulation of extracellular protease production to five Rap-Phr systems (RapC, F, F2, I1 and RapLike).

We found that the Spo0A phosphorelay and production of extracellular proteases are highly interconnected in Bt8741 through the functions of RapC, F and RapLike. Additionally, extracellular proteolytic activity (specifically the NprA protease) is regulated by the QS system NprR-NprRB (61), which is also involved in the modulation of the Spo0A phosphorelay (41, 42). Likewise, NprR also participates in the spreading phenotype of Bt8741 (A. Verdugo *et. al*, unpublished data), as well as RapC and RapLike. Because sporulation, extracellular protease production and spreading of Bt have evolved to be regulated and coordinated by multiple QS systems, these collective traits may be important in the life cycle of Bt and represent essential mechanisms for its ecology.

Mature Phr signaling peptides from Bs correspond to at least five residues located in the C-terminal end of the pro-Phr or in the middle of the sequence. Sequence analyses of mature Phrs in Bs have shown that a basic amino acid is found in the second position from the N-terminal end, and an alanine residue is necessary in the position before the cleavage site for Phr maturation (22, 62, 63). Our analysis of consensus putative mature Phr sequences showed that these characteristics are not maintained in mature Phr peptides of the *B cereus* group. This suggests that signaling peptides are processed differently in these bacteria, i.e., using different sets of extracellular proteases and peptidases that recognize distinct sequences. In Bt, the identity of a mature Phr has only been shown for Rap8-Phr8 from Bt-HD73. In this case, the active heptapeptide YAHGKDI is located in the C-terminal end from its exported sequence (46). RapI from Bt8741, ortholog protein to Rap8, is found in pherogroup 1, in which the consensus sequence HGKDI corresponds to the five residues in the C-terminal end from the exported sequence. This indicates that the consensus sequences determined in this study may not exactly predict the signaling peptide sequence, but they can direct their search in future studies.

We found that Rap-Phr systems in the *B. cereus* group have evolved into five pherogroups, each including Rap homologs from different species. This means that signaling peptides shared by more than one species, could mediate crosstalk or eavesdropping phenomena in nature, allowing the regulation of collective functions in response to interspecific signals as described for other gram-positive species (64, 65). The *B. cereus* group comprise bacteria with clinical and biotechnological relevance such as Ba, Bc, Bt, and other environmental and facultative species (38). We show that Rap-Phr QS systems in Bt are involved in the regulation of ecologically important collective traits, and our findings are highly relevant for further studies about the *B. cereus* group and contribute to the knowledge about its ecology. Understanding the regulatory processes for cell differentiation and specialization in these bacteria may enhance the use of biotechnologically-relevant species, or the strategies to control human pathogens, through the intervention of their collective functions at the molecular level. For instance, Ba and Bc are known for their pathogenic nature against mammals; therefore, elucidating the role of Rap-Phr systems in the production of virulence factors of these species such as anthrax toxin and capsule of Ba, or enterotoxins of Bc, could be of high relevance. Additionally, it is known that QS systems can be synthetically engineered (66, 67). As a result, Rap-Phr systems could be manipulated in order to enhance Bt survival, insect pathogenesis or cry protein production. This work serves as a starting point for the study of cell specialization of the *B. cereus* group bacteria.

## Materials and Methods

### Bacterial strains, media and culture conditions

*Bacillus thuringiensis* strain 8741 (Bt8741) (43), derived from Bt407 (Acc. No. NC_018877.1, 51), was used as host for the overexpression of Rap proteins. *Bacillus subtilis* strain 168 (Bs168) was used for the amplification of *rapA*. *Escherichia coli* strain TOP10 (69) was used for construction and cloning of overexpression plasmids before transforming into Bt8741. Luria-Bertani (LB) broth (10 g L^−1^ tryptone, 5 g L^−1^ yeast extract and 5 g L^−1^ NaCl) and Nutrient Agar (8 g L^−1^ nutrient broth, 15 g L^−1^ agar) were used at 30 °C for *Bacillus* cultures and at 37 °C for *E. coli* and 200 rpm for liquid cultures. Milk Agar was prepared using Nutrient Agar, supplemented with 5% skim milk (41). When needed, ampicillin (100 μg mL^−1^) or erythromycin (5 μg mL^−1^) was added to media. To induce expression from the *xylA* promoter in Bt8741, xylose was used to a final concentration of 20 mM (70), unless otherwise specified.

### Analysis of putative Spo0F-binding amino acids in Raps from Bt407

Based on the RapH residues involved in Spo0F binding in Bs168 (34) we determined the conservation of the corresponding residues in Raps from Bt407, in order to predict their capacity to bind to Spo0F. First, we analyzed the conservation of full-length Rap proteins from Bs168 and Bt8741 in comparison to RapH from Bs168. For this, we performed pairwise alignments of RapH amino acid sequence (NP_388565.2) with RapA (NP_389125.1), RapB (NP_391550.1), RapE (NP_390460.2), RapJ (NP_388164.1), RapD (NP_391519.1) from Bs168, and each of the eight Raps from Bt407 (AFV21721.1, AFV22194.1, AFV22088.1, AFV16731.1, AFV19251.1, AFV22208.1, AFV16776.1, AFV17466.1), using the BlastP tool (71). Then, all sequences were aligned together using MAFFT version 7 online service (72) with the G-INS-i iterative refinement method (73). Finally, we identified in the alignment the amino acids of Rap protein sequences that correspond to the residues of RapH that participate in binding and dephosphorylation of Spo0F.

### DNA manipulation

All primers used in this study are listed in Table S4. DNA was isolated from Bs168 and Bt8741 using the PureLink Genomic DNA Mini Kit (Invitrogen, Carlsbad CA, USA). QIAprep Spin Miniprep Kit (Qiagen, Germantown, MD, USA) was used routinely for plasmid extraction and purification. Oligonucleotides were designed for amplifying each Rap gene from Bt8741 genome or plasmids (Acc. No. NC_018877.1, NC_018883.1, NC_018886.1, NC_018879.1, NC_018878.1) and Bs168 genome (Acc. No. NC_000964.3), and synthesized as a commercial service (T4 Oligo, Irapuato, Mexico). PCR products and restriction reactions were purified using the PureLink Quick PCR Purification Kit (Invitrogen). When needed, PCR products were isolated from 0.8% agarose gels using the Zymoclean™ Gel DNA Recovery Kit (ZYMO Research, Irvine, CA, USA). Enzymes Dream Taq Master Mix, *HindIII*, *SalI* (Thermo Scientific, Waltham, MA, USA), *PstI* and T4 DNA Ligase (New England Biolabs Inc., Ipswich, MA, USA) were used as recommended by the manufacturer.

### Construction of Rap-overexpression Bt8741 strains

All strains and plasmids used in this study are listed in Table S1. For the construction of the overexpression plasmid pHT315-P_*xylA*_, the regulatory region of the xylose operon, including the *xylA* promoter (P_*xylA*_) and the repressor gene *xylR*, were amplified by PCR from Bs168 genome using primers GG1 and GG2 (Table S4). This PCR product was inserted into the *Hin*dIII and *Pst*I sites of pHT315 plasmid (74), and colonies were PCR checked using primers DS16 and DS17 (Table S4). The resulting plasmid pHT315-P_*xylA*_ was transformed into *E. coli* Top10 competent cells. Then, this plasmid was used for the inducible overexpression of Rap proteins with xylose in Bt8741. For this, *rap* genes encoded in the genome of Bt8741 (*rapC*, *rapK*, *rapF*, *rapF1*, *rapF2*, *rapI*, *rapI1* and *rapLike*, 47) and *rapA* from Bs168 (RapA_Bs_, 32) were amplified using the corresponding primers pairs listed in Table S4, and inserted in-frame between the *PstI* and *SalI* sites of pHT315-P_*xylA*_. Nine overexpression plasmids, one for each Rap protein, were transformed into *E. coli* Top10 competent cells. All plasmids were then transformed into Bt8741 electrocompetent cells, using the protocol described in previous studies (41), generating nine Bt8741 strains for the overexpression each Rap protein. Additionally, we transformed Bt8741 with the pHT315-P_*xylA*_ (without a *rap* gene), and the resulting strain was used as control strain throughout the Rap induction experiments. The complete sequence of pHT315-P_*xylA*_’*rapI* was verified by Illumina sequencing (MGH DNA Core, Cambridge, MA, USA), and the rest of the P_*xylA*_’*rap* constructions were verified by Sanger sequencing (Unidad de Servicios Genómicos, LANGEBIO-CINVESTAV, Irapuato, Mexico) using primers GG26 and DS17 (Table S4).

### Sporulation efficiency

We assessed the effect of the overexpression of Rap proteins on sporulation efficiency in Bt8741. Preinoculums were prepared by picking a single colony of each strain into 5 mL of liquid media and grown overnight. Then, 1 mL of preinoculum was centrifuged, washed and suspended in 1 mL of sterile PBS. Glass culture tubes (25 mm diameter) with 5 mL of LB with erythromycin were inoculated with 50 μL (1% v/v) of preinoculum containing ≈10^7^ cfu ml^−1^ and incubated for 72 h. All strains were cultured in triplicate, in LB with and without the addition of xylose. To determine growth and sporulation, total and thermoresistant CFU were calculated by plating 10-fold serial dilutions in nutrient agar. For thermoresistant CFU, samples of 100 μL were incubated at 80 °C for 20 min prior to diluting and plating. Sporulation efficiency was calculated as the percentage of thermoresistant CFU in total CFU.

### Biofilm formation assay

We evaluated the effect of the overexpression of Rap proteins on the capacity of Bt874 to form biofilms. For this assay, we used 13 × 100 mm glass tubes with 3 mL Nutrient Broth + erythromycin, with and without the addition of xylose to a final concentration of 2 mM. Three μL of preinoculum was added in triplicates, and the inoculated tubes were incubated without agitation at 31 °C ± 1 °C for 48 hours. The culture media was then removed with a syringe with needle. The biofilm and ring attached to the wall of the tube, composed of cells from the biofilm, were suspended in 1.5 mL of sterile PBS and the optical density (OD_600_) was measured. The OD_600_ was also measured from the removed liquid media to address planktonic growth. At least 5 replicates of each treatment were performed.

### Extracellular proteolytic activity assay

To evaluate the effect of Rap overexpression in extracellular proteolytic activity of Bt8741, 2 μL of preinoculums of each Rap-overexpression strain, prepared as described above, were spotted in triplicate on milk agar with and without the addition of xylose. The hydrolysis halo area was measured after 24 h of incubation using the Image Lab™ Software (BIORAD). To correct for differences in colony growth, we subtracted the colony area.

### Spreading phenotype assay

The spreading phenotype of Rap-overexpression Bt8741 variants was followed in colonies spotted on agar. For this assay, we used diluted nutrient agar (NA) (0.8 g L^−1^ Nutrient broth, 1.5 g L^−1^ agar) with erythromycin and with or without the addition of xylose. Plates were air-dried inside a biological hood for 60 minutes prior to inoculation. Then, 5 μL of preinoculum cultures were spotted in the center of the plate, dried for 5 minutes and incubated at 30 °C for 14 days. The inoculated agar plates were photographed at days 1, 3, 5, 7 and 14, using a gel documentation system (Gel Doc™ XR+, BIORAD). Colony area was measured using the Image Lab™ Software (BIORAD) and radial growth was calculated. For normalization of radial dispersion, we subtracted from all observations the colony radius at day 1, which corresponds to the inoculated droplet area. Three replicates of each treatment were performed.

### Phylogenetic Analysis

To reconstruct the phylogeny of Rap proteins in the *B. cereus* group, we first selected one representative strain of each species from NCBI GenBank, including *Bacillus cereus* ATCC14579 (Accession NC_004722.1), *Bacillus anthracis* A0248 (NC_012659.1), *Bacillus thuringiensis* 407 (NC_018877.1), *Bacillus mycoides* ATCC6442 (NZ_CP009692.1), *Bacillus pseudomycoides* DMS12442 (NZ_CM000745.1) and *Bacillus cytotoxicus* NVH391-98 (NC_009674.1). Strains were selected based on the availability of a complete genome (as of July of 2018) and thus, *Bacillus weihenstephanensis* was excluded. We searched for Rap protein homologs in the selected genomes by querying the amino acid sequence of *B. subtilis* RapA (NP_389125.1) and each of the eight Rap sequences of *B. thuringiensis* 407: RapC (AFV21721.1), RapK (AFV22194.1), RapF (AFV22088.1), RapF1 (AFV16731.1), RapF2 (AFV19251.1), RapI (AFV22208.1), RapI1

(AFV16776.1) and RapLike (AFV17466.1). Homologs were searched using BLAST tool (71), the tBlastn tool and a local script designed for performing the blast search in an assembled database of the selected genomes. To ensure the identity of the Rap protein homologs, Blast hits were submitted manually to the Conserved Domain Search-NCBI tool (75) in order to determine if they presented the characteristic TPR-containing domain. Rap protein amino acid sequences were aligned in MAFFT version 7 (72) using the G-INS-i iterative refinement method which incorporates pairwise alignment algorithms (73). RapA and RapH from Bs168 were also included as outgroups for the phylogenetic reconstruction. The selection of the best substitution evolutionary model (JTT+G+I+F) was made using the Smart Model Selection with the Akaike Information Criterion in PhyML 3.0 (76, 77), as well as the phylogeny reconstruction by the Maximum Likelihood method using 1000 bootstraps to support the phylogenetic prediction.

### Phr pro-peptide identification and pherogroup prediction

Additional to the identification of Rap homologs in the *B. cereus* group, we also analyzed the putative *phr* genes, which code for pro-Phr, the precursor of the quorum sensing signal peptide. For this, we performed a manual search targeting open reading frames (ORFs) between 30 and 100 amino acids of length, downstream from the *rap* gene sequences. When present, each Phr amino acid sequence was analyzed for the presence of a signal peptide for secretion and a cleavage site using SignalP4.1 (78). The putative mature signaling peptide (mature Phr) and pherogroup prediction were performed from the exported Phr amino acid sequences (pro-Phr). For this, Phrs corresponding to Rap proteins from different clades of the phylogenetic reconstruction were analyzed separately. The amino acid sequences of the pro-Phr from each clade were aligned using ClustalW (79). Pherogroups were identified by manually, by modifying the groups of aligned Phrs and looking for consensus sequences in the alignments. For better identification of consensus sequences, sequence Logos were created for each pherogroup using the Seq2Logo 2.0 online service (80).

### Statistics

All the statistical analyses were performed using GraphPad Prism version 7.0a. Data obtained from the extracellular proteolytic activity assay, spreading (at day 7) and biofilm formation were analyzed with multiple *t*-tests to search for differences between not induced and induced Rap protein overexpression conditions of each strain. Significance of 0.05 was used in all statistical tests.

## Supporting information

supplemental material is available.

## Acknowledgements

Authors thank all members of the Microbe-Plant Interactions group at CIDEA for their comments and suggestions. We gratefully acknowledge Gabriela Olmedo-Álvarez for her contributions in analyses of phylogenetic data, suggestions throughout the development of the experimental work and critical review of the manuscript. Bernardo Aguilar and Enrique Hurtado contributed with technical assistance in the phylogenetic analyses. GG Received a scholarship 636324 from Conacyt. This work was partially funded by Conacyt (Fomix 267837 and Fordecyt 296368) for MT.

## Notes

#### Summary of Updates

Minor corrections in the text and captions. Figures updated with more visible labels. References in tables corrected to match overall number formatting.

https://github.com/gabyga16/rap_phylogenetics

