## supplemental material is available. for "Rap-protein paralogs of *B. thuringiensis*: a multifunctional and redundant regulatory repertoire for the control of collective functions"

### Supplementary Figures.

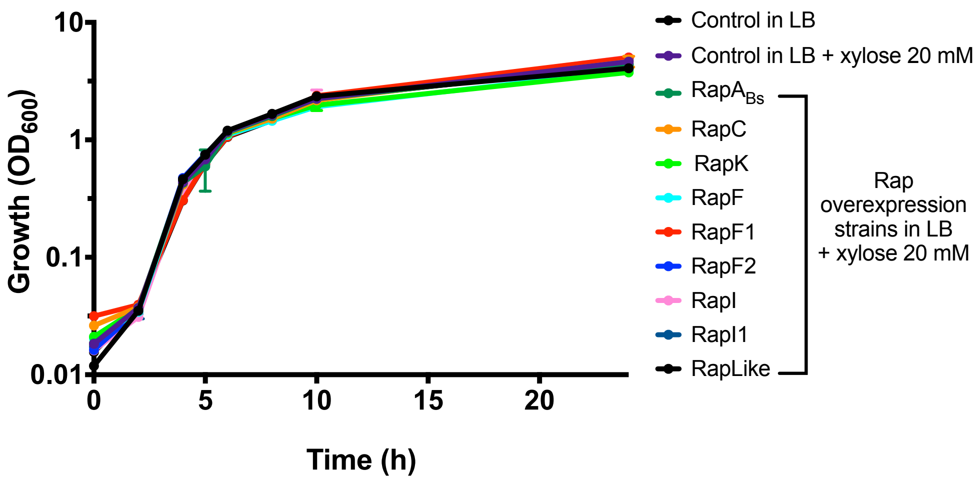


**Figure S1.** Growth curves of Rap-overexpression strains. Pre-inoculums of the control strain, and strains carrying plasmids for Rap overexpression, were washed and used to inoculate 30 ml of fresh media to a starting $DO_{600}$ ≈0.03, in triplicate 125 ml flasks. Cultures were incubated at 30 ºC with shaking, following $DO_{600}$ for 24 h. Each data point represents average of three replicates ± SD.

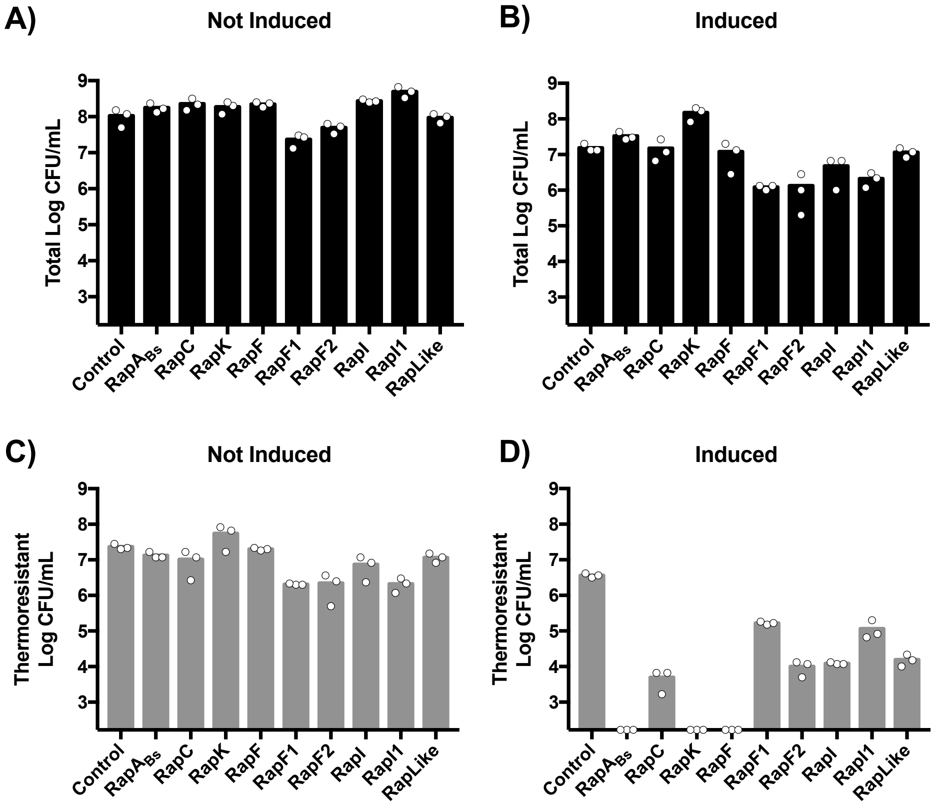


**Figure S2.** Effect of the induction of Rap overexpression in viability and sporulation of Bt8741. Total and thermoresistant CFU counts were obtained from triplicate 72 h cultures of control and Rap-overexpression strains, from LB with and without the addition of 20 mM xylose. Columns indicate average of three replicates; dots indicate individual data. Y-axis lower limit is adjusted to the limit of detection of the assay.

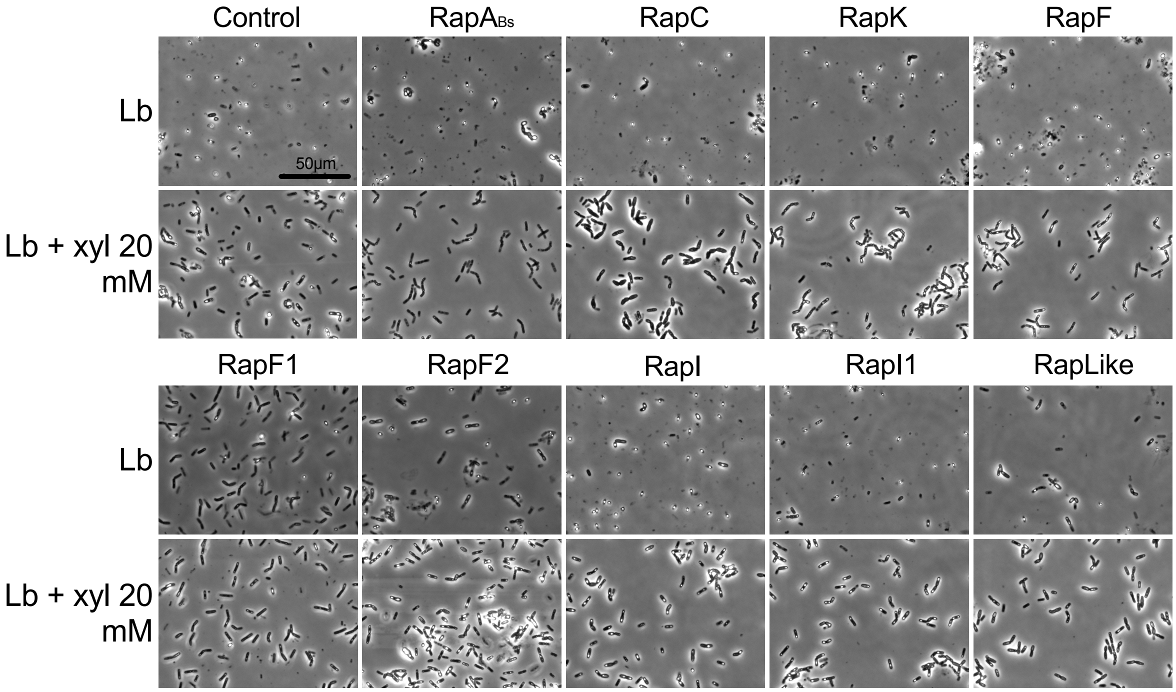


**Figure S3.** Cell morphology of control strain and Rap-overexpression strains. Samples were taken from induced cultures at 72 h. Phase contrast microscopy 63X + 1.8X magnification.

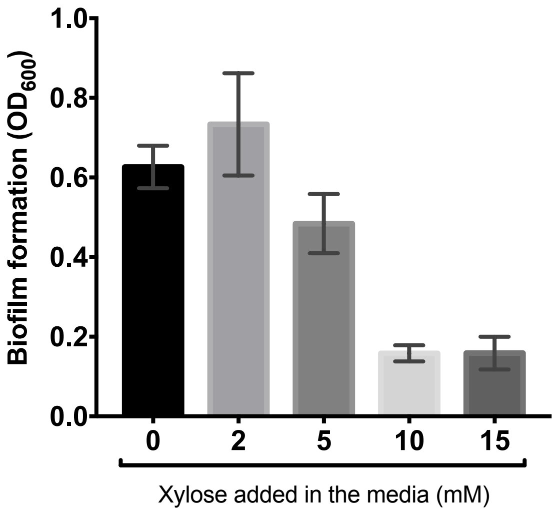


**Figure S4.** Biofilm formation of Bt8741 is inhibited by xylose addition. Three µl of preinoculum of the control strain, were used to inoculate 3 mL of nutrient broth with different concentrations of xylose. After 48 h, cells from the biofilm were recovered, suspended in 1.5 ml of PBS, and $OD_{600}$ was measured. Columns indicate average of three replicates ± SD.

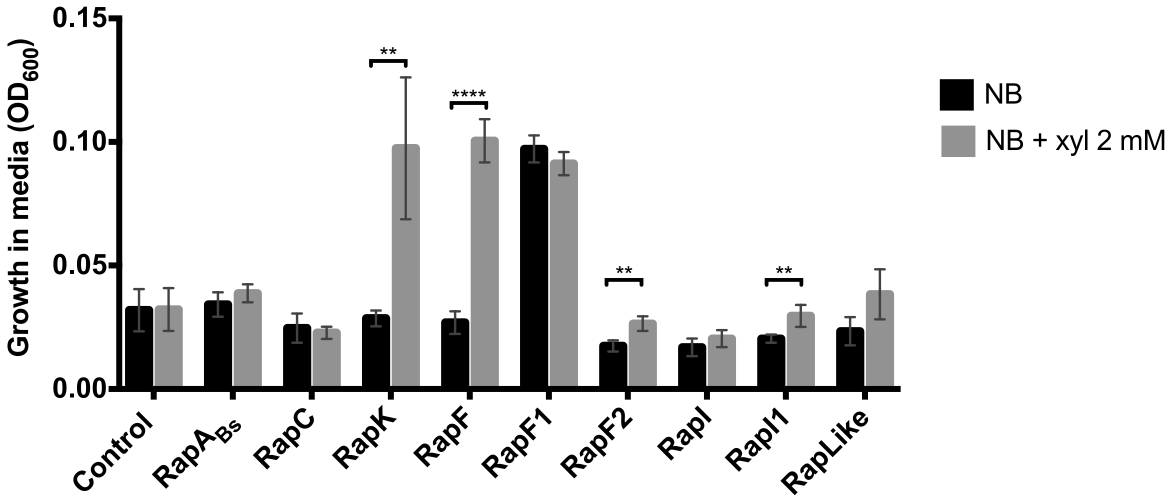


**Figure S5.** Planktonic growth of control and Rap-overexpression strains, during the assays for biofilm formation. $DO_{600}$ was measured from the liquid media after 48 h. Columns indicate average of five individual measurements ± SD. **, $p<0.005$; ****, $p<0.0001$.

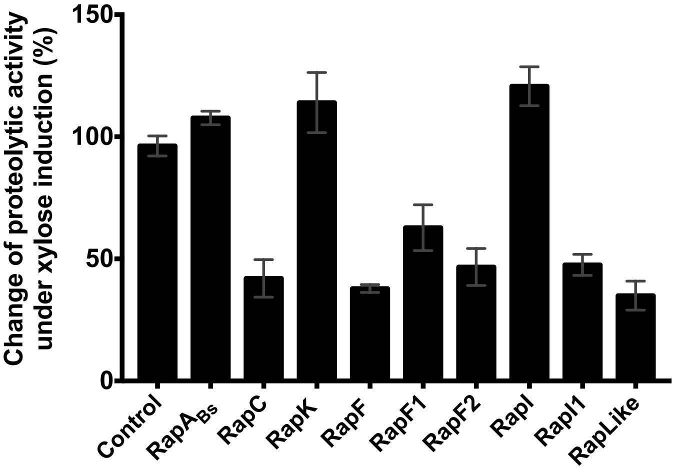


**Figure S6.** Inhibition of proteolytic activity by Rap proteins. The effect of Rap overexpression on extracellular proteolytic activity was calculated as percentage of proteolysis activity, in induced media, with respect to non-induced media. Columns indicate average of triplicate measurements ± SD.

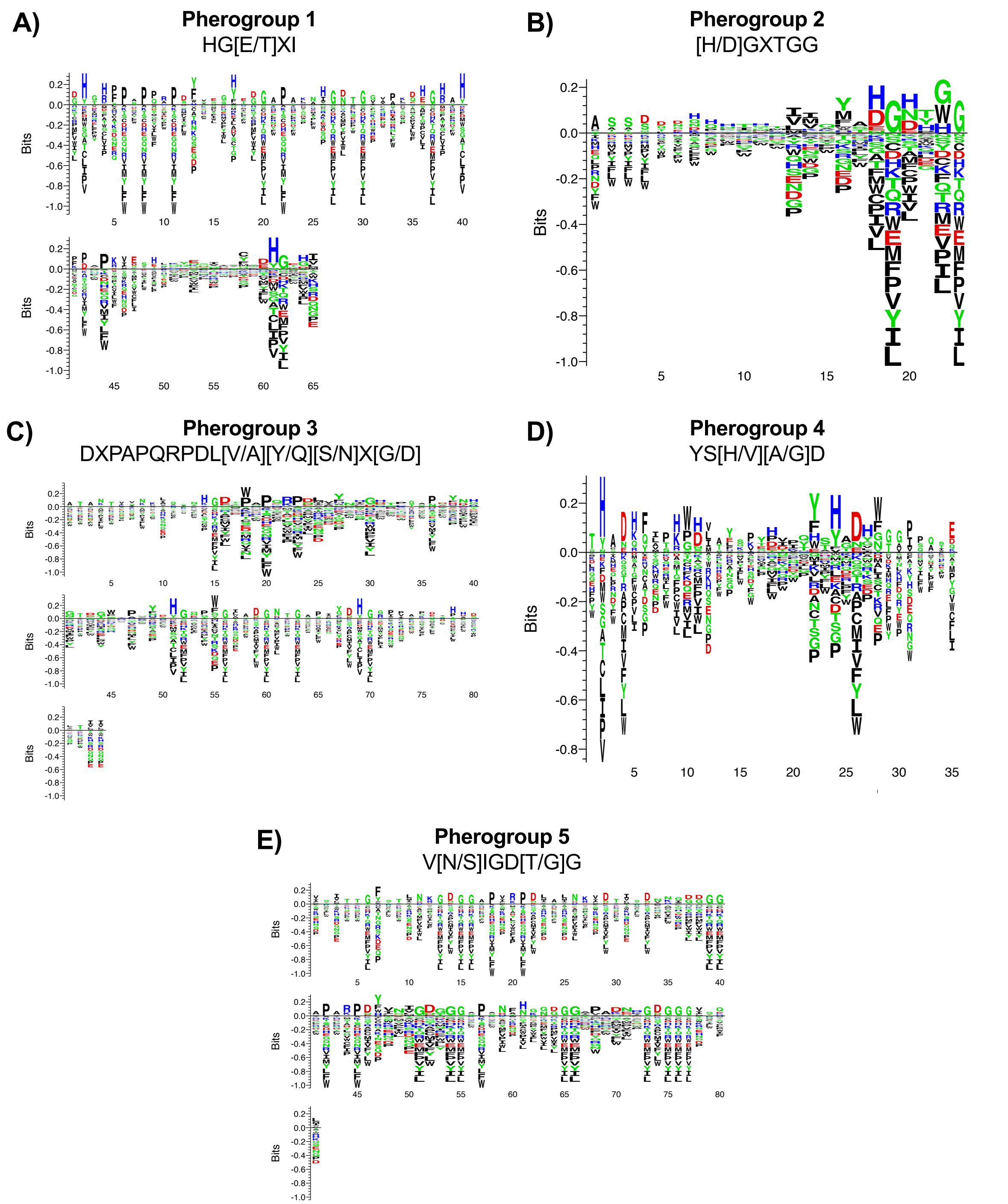


**Figure S7.** Phr pherogroups from species from the *B. cereus* group. Multiple alignments of exported Phr sequences were used for obtaining LOGOS, using the Seq2Logo 2.0 online tool (http://www.cbs.dtu.dk/biotools/Seq2Logo/). Consensus sequences are shown for each Logo.

**Supplementary Tables.**

**Table S1.** Strains and plasmids used in this study

| Strain | Description | Reference |
|---|---|---|
| *Bacillus thuringiensis* 8741 | Wild type | (43) |
| *Bacillus subtilis* 168 | Wild type | (32) |
| *Escherichia coli* TOP10 | Strain used for construction and cloning of overexpression plasmids | (69) |

| Plasmid | Description | Reference |
|---|---|---|
| pHT315 | Cloning vector for Bt ≈15 copies per cell | (74) |
| pHT315- $P_{xylA}$ | Xylose inducible overexpression plasmid | This study |
| pHT315- $P_{xylA}$'*rapC* | Plasmid for overexpression of RapC from Bt8741 | This study |
| pHT315- $P_{xylA}$'*rapK* | Plasmid for overexpression of RapK from Bt8741 | This study |
| pHT315- $P_{xylA}$'*rapF* | Plasmid for overexpression of RapF from Bt8741 | This study |
| pHT315- $P_{xylA}$'*rapF1* | Plasmid for overexpression of RapF1 from Bt8741 | This study |
| pHT315- $P_{xylA}$'*rapF2* | Plasmid for overexpression of RapF2 from Bt8741 | This study |
| pHT315- $P_{xylA}$'*rapI* | Plasmid for overexpression of RapI from Bt8741 | This study |
| pHT315- $P_{xylA}$'*rapI1* | Plasmid for overexpression of RapI1 from Bt8741 | This study |
| pHT315- $P_{xylA}$'*rapLike* | Plasmid for overexpression of RapLike from Bt8741 | This study |
| pHT315- $P_{xylA}$'*rapA* | Plasmid for overexpression of RapA from Bs168 | This study |

**Table S2**. Rap-Phr systems found in representative strains of the *B. cereus* group.

| | | Rap | | Phr |
|---|---|---|---|---|
| **Specie** | **Location** | **Phylogeny identifier** | **NCBI Acc. No.** | **NCBI Acc. No.** |
| ***Bacillus subtilis* 168** | Chromosome | BsRapA | NP_389125.1 | NP_389126.1 |
| | | BsRapH | NP_388565.2 | YP_003097693.1 |
| ***Bacillus cereus* ATCC 14579** | Chromosome | Bc3518 | NP_833251.1 | NP_833252.1 |
| | | Bc1026 | AAP08013.1 | No Phr |
| | | Bc3501 | NP_833234.1 | NP_833233.1 |
| | | Bc2147 | NP_831913.1 | NP_831914.1 |
| | | Bc0986 | AAP07973.1 | EEL12896.1*** |
| ***Bacillus anthracis* str. A0248** | pXO1 | Ba29315 | ACQ50983.1 | ACQ50984.1 |
| | Chromosome | Ba12985 | ACQ49614.1 | ACQ47217.1 |
| | | Ba05875 | ACQ48214.1 | ACQ48469.1 |
| | | Ba17875 | WP_001102720.1** | ACQ48710.1 |
| | | Ba16555 | ACQ46490.1 | ACQ46858.1 |
| | | Ba18790 | ACQ46549.1 | ACQ48053.1 |
| ***Bacillus thuringiensis* 407** | pBTB_6p | BtRapK | AFV22194.1 | EEM24826.1 |
| | pBTB_9p | BtRapI | AFV22208.1 | AFV22209.1 |
| | pBTB_78p | BtRapF | AFV22088.1 | AFV22087.1 |
| | pBTB_502p | BtRapC | AFV21721.1 | AFV21722.1 |
| | Chromosome | BtRapF1 | AFV16731.1 | AFV16732.1 |
| | | BtRapF2 | AFV19251.1 | AFV19252.1 |
| | | BtRapI1 | AFV16776.1 | AFV16777.1 |
| | | BtRapLike | AFV17466.1 | AFV17465.1 |
| ***Bacillus mycoides* ATCC 6442** | pBMX_1 | Bm01680 | WP_003192987.1** | WP_003192985.1** |
| | Chromosome | Bm14240 | CP009692.1* | WP_002165311.1** |
| | | Bm28480 | WP_003193466.1** | WP_003193464.1** |
| | | Bm15830 | WP_003190346.1** | WP_016127268.1** |
| | | Bm25900 | WP_003187761.1** | WP_002140709.1** |

| Specie | Location | Rap | | Phr |
|---|---|---|---|---|
| | | Phylogeny identifier | NCBI Acc. No. | NCBI Acc. No. |
| ***Bacillus cytotoxicus* NVH 391-98** | Chromosome | Bcyt11595 | ABS22527.1 | ABS22528.1 |
| | | Bcyt18870 | ABS23932.1 | ABS23933.1 |
| | | Bcyt04205 | ABS21090.1 | SCL85822.1*** |
| | | Bcyt20060 | ABS24169.1 | ABS24170.1 |
| | | Bcyt05405 | ABS21319.1 | SCN32895.1*** |
| | | Bcyt02700 | ABS20819.1 | SCN30732.1*** |
| | | Bcyt05320 | ABS21305.1 | SCN32816.1*** |
| | | Bcyt09890 | ABS22207.1 | ABS22206.1 |
| ***Bacillus pseudomycoides* DMS 12442** | Chromosome | Bps04625 | EEM18248.1 | EEM18249.1 |
| | | Bps28285 | EEM13381.1 | Not annotated |
| | | Bps11660 | EEM16725.1 | EEM16724.1 |
| | | Bps15675 | EEM15767.1 | EEM15766.1 |
| | | Bps05775 | EEM17495.1 | AIK39315.1*** |
| | | Bps17975 | EEM15413.1 | EEM15414.1 |
| | | Bps14840 | EEM16052.1 | WP_033799691.1** |
| | | Bps24285 | EEM14137.1 | AIK40762.1*** |

* Truncated protein.
** Access number corresponding to an NCBI reference sequence.
*** Access number corresponding to a protein in a different strain of the same species.

**Table S3.** Pherogroups of signaling Phr peptides. Consensus sequences are highlighted in gray.

| Pherogroup | Phylogeny Identifier | Predicted signaling peptide* | Signaling peptide* | Exported sequence* |
|---|---|---|---|---|
|  | Bc2147 | Yes | MKKAILLMMMLSAVFTFGLTNA | ----------------------------------------KDIP------QNEAIEVFMDHGEHI |
|  | Ba18790 | No | MKKIKITLMGLIGIAVLSFGLNSPSIEKSA | ------------------------------------------AP----TLKQENIISYSEHGTGI |
|  | **BtRapI** | Yes | MMKKFSLILIGVACTTGIFFSQFNNSIQTHDA | -------------------------------------------------KEKNDIIQQYAHGKDI |
| **1** | Bm15830 | No | MKKTIGKIAVGIVFTGIAAFGLNAGFTNQA | GHTHFPSPPAPSFSEGHTGGAPALNIGDTGGAPKQEGHAHFPAPKVESHGNTGGAPANSEHGTGI |
|  | Bps11660 | No | MKKVLLTVMIFTGILSFGVANQSIE | ------------------------------------------EP------SAKANLLCLDHGETI |
|  | Bcyt09890 | Yes | MKKIKIALLGLMSVAVVSFGLNSPITKDNA | ------------------------------------------SS----ELKYETVSYMSEHGGGI |
|  | Bcyt18870 | Yes | MKKVLMGFVSFAAVLTIGSFAESYTSQA | DHGRPPAPQRPDYVSSYED----KAHGNTGVVALDHGR--PPAP------AYEYGDVYTDHGEHI |
|  | **BtRapF** | Yes | MKMKKTLISLMGIVTILTFTIGLSNPGEVQK | --------FIQTKVAASEGDYGG |
| | Ba05875 | Yes | MKKMVIKLLGIVTVFTLTLGVYS | ---STNHQKSPTLMKAEHGDYGG |
| | **BtRapF1** | No | MKRIISSSIGLIITVILLNGVNVST | ---DQLKVNTTSVIQYTHGEPWG |
|  | Ba29315 | Yes | MKKVMFSLIGLTAVFTFMFNASNVTDTQKA | -------LSEDKVVQYAHGHTGG |
|  | Bcyt11595 | No | MRKVIFSLIGITAAFTFMFNSSHVTDTQKA | -------LNEDTIVQYAHGHTGG |
| | Bcyt05405 | Yes | MKMKKTVLSLLGVLTVLTLTFG | ASSSLDVQQTVEIIAYTHGHTG- |
| **2** | Bps28285 | Yes | MKFLKLSLKRFIAVGILAAGISSFGFNAQPTDVQQ | --------ASSAVVSYAHGNTG- |
|  | Bcyt05320 | Yes | MKKMIGSIMGSMMVLTLTFGIHTLVEKPQYA | ----DGNHHKMMLIQYADGNHH- |
|  | Bps05775 | No | MKKVILGLMGIVTILTLTLGVYTTVD | -----EQQTTTLVKTMADGNTGG |
|  | **BtRapK** | Yes | MKKTILTLMGIITVFTLTLSNINTPKEN | -----KDPSIQKIMLMSDGNTGG |
|  | Bps24285 | Yes | MKKVLLGMLLITFLAVSTHTSVGT | ------QTVTNLIQSLADGDTG- |
|  | Bcyt02700 | Yes | MKMKKVIISLMGIMTALTLTFGVLDSKDTHQA | ---------IHTVKQMSDGHTGG |

| Pherogroup | Phylogeny Identifier | Predicted signaling peptide* | Signaling peptide* | Exported sequence* |
|---|---|---|---|---|
| 3 | Ba17875 | Yes | MIKKISSVFLGLSVFGILATGIHSSFTYQAGHA | ---------------DLPAPQRPDLVQSVDASK---DYNST-TTAEKAL----------------------------------- |
|  | **BtRapF2** | Yes | MIKKISSIVLGLSVLSIVSIGLNSSFTYQAGHA | ---------------DFPAPQRPDLVQSIDVSK---DYNST-TTEKAL------------------------------------ |
|  | Bm14240 | Yes | MKKLSLMVMSLAVTGITVFTINTTPKIQQT | ATANTVVNKLQSTHGEGWSPTRPELVYLEGRQL--------------------------------------------------- |
| | Bm01680 | No | MKKIGLTIAGLAVLGITSLGLGHTNEIA | ---KTKQKLIAVNIGDTWSPQRPDLAYSHGHTGGSPEYTHG-EGWSPSYTHGEPWGIADGNTGAPAYDHGRPPAPTNTHGETII |
|  | Bm28480 | Yes | MKKLKTALLLSVTAFSFLAMAEG | ---------------DTPAPQKPGLAYSHGHTGGSPEYTHG-EGWSPSYTHGEPWGIADGNTGSPAYDHGRPPIPQKPDLV--- |
|  | Bps17975 | No | MKKIGLTITGLALAGIMTFGVSNNPGFSSL | ------------NKGDTPAPQRPDLALNIGDTP-APADNGG-DTPAPAYDHGQPWGLADGNTGASIGDHGGIVAQDTHADL--- |
|  | Bps15675 | Yes | MKKLKAVLLLTVTSFGLLSMAEG | ---------------DTPAPQRPDLVYNIGDTP-APAYNRG-DGGSPSNAHGNGGGI---------------VAKEI------- |
|  | Bcyt20060 | Yes | VMKKLCSIVLSLAIIGIVSFGLNSSIEILQS | ------------AHGDYPAPQRTNLVYNLGNTLTSNNSDGNTGGIIANYEHGQTF----------------------------- |
| 4 | Bc3518 | No | MKKISLLLSGIAFIGVLTVGIFQFSKTDQL | ------ATHGH-----YPAPSYSVGDYGGAPPQI- |
|  | Bc0986 | Yes | MKKLVLATLGLVIALSFSNNTGDTQH | ------------ALKKEEIIQYSHADHF------- |
|  | Ba16555 | Yes | MKKISLLLSGIAFIGILTVGISPLSKA | ---DQFATHGH-----YPVPTYSVGDHGGVPQQT- |
|  | **BtRapLike** | Yes | MKKISLAILTFTCILAFGFNNFTESQQA | ----KQLPKWD------TDKQETHADL-------- |
|  | **BtRapC** | Yes | MKKFKIALLGFVSVAVLSLGLNSGTETQKA | ---------SSVTESPTHVIYYSHADVW------- |
| | Bps04625 | Yes | MKKLGVITVGLSVIGILSLGLNHFAETQQA | THADHFISKPDLAYSVSDTSTYSYTDGNTGISASE |
|  | Bcyt04205 | Yes | MKMMKKFVFATLGLVMALAIGWNNEGDTQHA | -------------LQKEEIIQYSHADHF------- |
| 5 | **BtRapI1** | Yes | MKKFRLAIVGTALVGVLSIGFNSSFTNQA | -----------------------------------------------VNIGDTG-----------GAPADNKGDGG-VSQL |
|  | Bm25900 | Yes | MKKVGVIAKGLIFAGVLSFGFNFSTTTQQA | ---------------------------------VNIGDGGAPARPDYVSIGDGGGAPANFNKGDGGAPADNRGDGGGVSQL |
|  | Bps14840 | Yes | IKKILLTTIGCAFA | VSITTGFSTLNKGDGGAPVRPDLALNKVDTISDNKGDGGGAPARPDYVNIGDGGGSPSNAHGNGGGIVAKEI--------- |
|  | Bc3501 | No | MVEGLCNKLAQSGYGGTFIIIKPNENPYA | ------------------------VS------------EDTTMPA--KQIFY--------------------------- |
|  | Ba12985 | Yes | MKKIKKLLFSTIIVSTLACSMISLNVSQH | ----------SSF-----------NG------------GDTPAPT--KLIISGGVSIN--------------------- |
| | Bc1026 | - | | |

*Prediction of signal peptide and cleavage site was made with SignalP4.1.

**Table S4.** Oligonucleotides used in this study.

| Name | Sequence (5' to 3') | Description |
|---|---|---|
| GG1 | AAGAAGCTTCCATGTCACTATTGCTTCAG | Fwd $P_{xylA}$ |
| GG2 | CTGCTGCAGAGATTGAGCCATGTGATTTC | Rev $P_{xylA}$ |
| GG3 | CTGCTGCAGGACGTTCAAACAAAAAGTAATG | Fwd *rapK* |
| GG4 | GTCGTCGACTTCCCATTAATGTTAGGATTG | Rev *rapK* |
| GG5 | CTGCTGCAGCAACTAGGAAACGAACAAATTAC | Fwd *rapI* |
| GG6 | GTCGTCGACTTCATCATTTCAATGACTCC | Rev *rapI* |
| GG7 | CTGCTGCAGACAGCAACCAGTAATGAGAAG | Fwd *rapF* |
| GG8 | GTCGTCGACGTTTTCTTCATTTTAATGCCTC | Rev *rapF* |
| GG9 | CTGCTGCAGAACACTAAGTTTTTGACCCAAG | Fwd *rapC* |
| GG10 | GTCGTCGACTTTTCATCACTGTAACGCTC | Rev *rapC* |
| GG11 | CTGCTGCAGAGTGTACATGTAATAAAAAAGGAAG | Fwd *rapF1* |
| GG12 | GTCGTCGACCAATTGAACTGCTGATGATC | Rev *rapF1* |
| GG13 | CTGCTGCAGAACGTTCAACTACAAGGTAATG | Fwd *rapF2* |
| GG14 | GTCGTCGACTATCATTTTAATGCCCCTTTC | Rev *rapF2* |
| GG15 | CTGCTGCAGGGAGCAGATGTAGTAACGC | Fwd *rapI1* |
| GG16 | GTCGTCGACATAGCCAAACGAAATTTCTTC | Rev *rapI1* |
| GG17 | CTGCTGCAGTTATTAAAAGGGCATGAACAG | Fwd *raplike* |
| GG18 | GTCGTCGACTCATTATCTTAGTGCCTCCTTAG | Rev *raplike* |
| GG19 | CTGCTGCAGAGGATGAAGCAGACGATTC | Fwd *rapA* |
| GG20 | GTCGTCGACAACAAACCTGACATCCATTTAG | Rev *rapA* |
| GG26 | AACATAGTACATAGCGAATCTTC | Fwd $P_{xylA}$ |
| DS16 | CAGGCTTTACACTTTATGC | Fwd pHT315 |
| DS17 | CGATTAAGTTGGGTAACG | Rev pHT315 |

Underlined sequence indicates restriction site; Fwd, forward primer; Rev, reverse primer.
